# A PARP2-specific active site α-helix melts to permit DNA damage-induced enzymatic activation

**DOI:** 10.1101/2024.05.20.594972

**Authors:** Emily S. Smith-Pillet, Ramya Billur, Marie-France Langelier, Tanaji T. Talele, John M. Pascal, Ben E. Black

**Affiliations:** Department of Biochemistry and Biophysics, Penn Center for Genome Integrity, Epigenetics Institute; Graduate Program in Biochemistry, Biophysics, Chemical Biology Perelman School of Medicine, University of Pennsylvania, Philadelphia, PA 19140-6059 USA; Department of Pharmaceutical Sciences, College of Pharmacy and Health Sciences, St. John’s University, Queens, NY 11439 USA; Département de Biochimie et Médecine Moléculaire, Université de Montréal, Montréal (Québec), H3C 3J7 Canada

## Abstract

PARP1 and PARP2 recognize DNA breaks immediately upon their formation, generate a burst of local PARylation to signal their location, and are co-targeted by all current FDA-approved forms of PARP inhibitors (PARPi) used in the cancer clinic. Recent evidence indicates that the same PARPi molecules impact PARP2 differently from PARP1, raising the possibility that allosteric activation may also differ. We find that unlike for PARP1, destabilization of the autoinhibitory domain of PARP2 is insufficient for DNA damage-induced catalytic activation. Rather, PARP2 activation requires further unfolding of an active site α-helix absent in PARP1. Only one clinical PARPi, Olaparib, stabilizes the PARP2 active site α-helix, representing a structural feature with the potential to discriminate small molecule inhibitors. Collectively, our findings reveal unanticipated differences in local structure and changes in activation-coupled backbone dynamics between PARP1 and PARP2.

## Introduction

PARP1 and PARP2 are related enzymes that both coordinate DNA repair and transcriptional regulation by catalyzing covalent attachment of poly(ADP-ribose) (PAR) to target molecules^1,2^. Both rapidly engage DNA breaks within seconds of their formation^3^ and immediately generate linear and branched chains of PAR, built from substrate NAD^+^, both in the form of automodification and modification of chromatin components, including histone H3Ser10 in nucleosomes proximal to the break^4–6^. Both enzymes are autoinhibited when unbound to a DNA break, but autoinhibition is relieved through pronounced destabilization of their autoinhibitory “helical domains” (HDs) that permits NAD^+^ access to the catalytic centers of the enzymes^7^. Additionally, the co-factor Histone PARylation Factor 1 (HPF1) confers rapid initiation by both PARP1 and PARP2 of serine modification on target proteins^4,8,9^. Both PARP1 and PARP2 are targeted by the four FDA-approved PARP inhibitors (PARPi) used in the cancer clinic^10^. Genetic evidence for having at least partially overlapping roles is exemplified by mouse experiments wherein either individual knockout produces viable animals, albeit sensitive to genotoxic treatments, but the double knockout (PARP1^-/-^; PARP2^-/-^) is inviable^11^.

Despite these indicators of similar function, regulation, and inhibition, there are marked differences between the two enzymes. The most obvious is in the N-terminal portions of the enzymes. PARP1 has Zn^2+^-binding domains, F1-F3, that are largely responsible for recognizing DNA breaks, as well as a BRCA1 C-terminal (BRCT) domain that are completely absent in PARP2 (Fig. 1A). In PARP2, DNA break recognition is mediated by the WGR domain (33% identical to the same domain in PARP1)^12,13^. The different DNA binding domain compositions lead to substantial changes in affinity for various types of DNA damage lesions^12^, and retention on DNA lesions is more pronounced for PARP2 than for PARP1^14^. In terms of allosteric communication through the enzymes, there is an indication that reverse allostery (i.e. from the NAD^+^ site back to impact the release or retention of PARP1 or PARP2 on a DNA break) occurs differently in PARP1 versus PARP2, with different outcomes from the same PARPi molecules on the two enzymes^15–17^. In PARP1, this involves changes in the local backbone dynamics of the allosteric HD^15^, whereas in PARP2, it appears to be explained primarily through relatively subtle structural movements within the HD^16^. With all of this, we predicted that despite some overall similarity^7^, PARP2 would employ unique mechanisms upon activation while engaging a DNA break.

**Figure 1:**
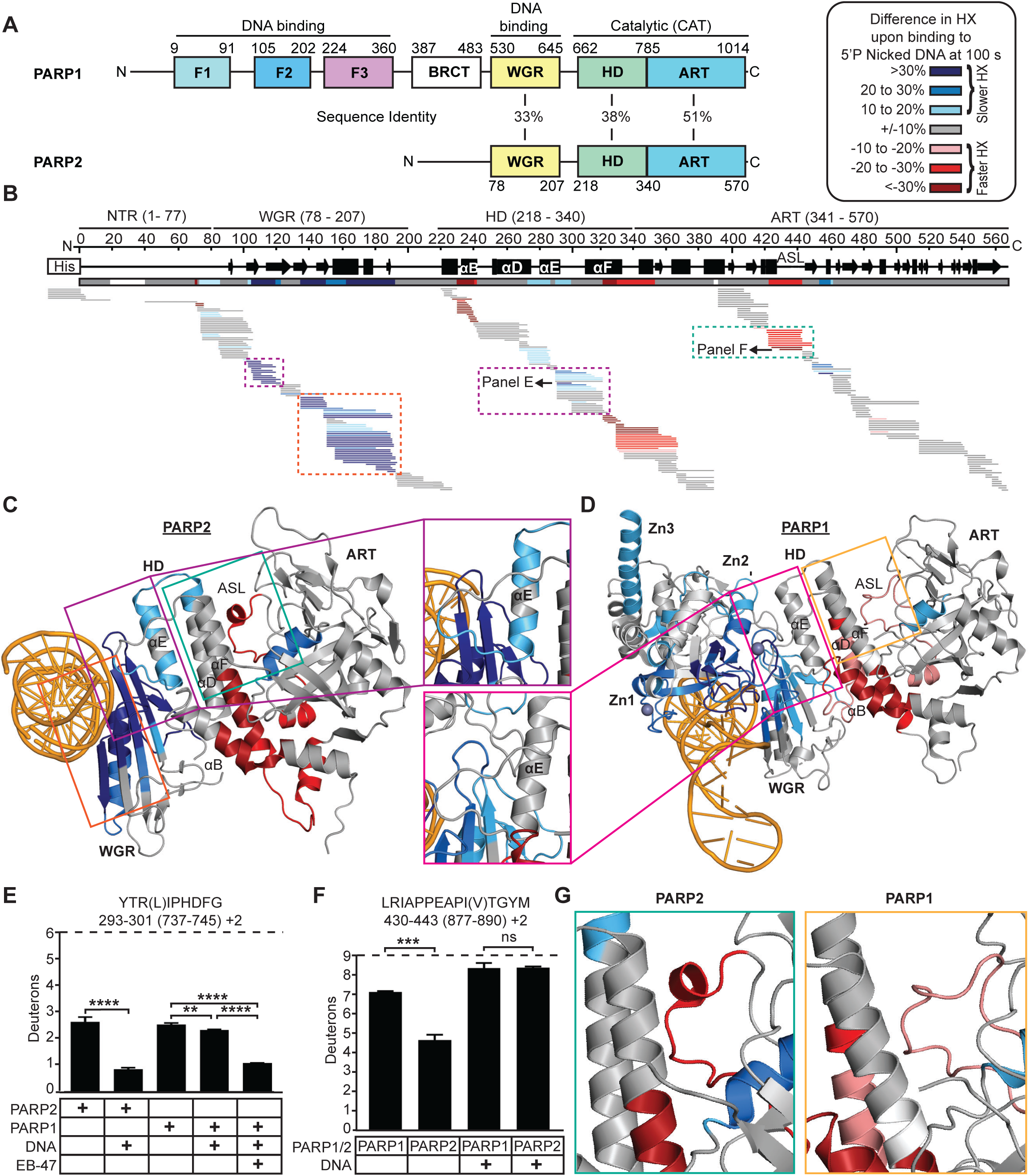
PARP2 exhibits local HX protection and deprotection from DNA break induced activation not observed in PARP1. (A) Domain architecture of PARP1 and PARP2 and the percent sequence identity between each shared domain is displayed. (B) HX difference plot between WT PARP2 and WT PARP2 complexed with 5’P nicked DNA at 100 s. Horizontal bar represent PARP2 peptides. White gaps represent the missing coverage. The color key of HX differences is shown next to panel A. (C) Consensus HX data from panel B mapped on the cryo-EM structure of PARP2 complexed with DNA and HPF1 (PDB 6X0L) with HPF1 removed for clarity. (D) Consensus HX data of PARP1 upon binding to DNA mapped on the crystal structure of PARP1 on DNA damage (PDB 4DQY: F3, WGR, HD, ART) combined with the NMR structure of PARP1 on DNA damage (PDB 2N8A: F1, F2, DNA). (E) HX at 100 s of representative peptides in the C-terminus of αE of PARP2 or PARP1 with the indicated condition (DNA and/or EB-47). PARP1 specific residues are shown in parenthesis. Each bar represents the average value from three replicates, with the error bars representing the SD. * and **** indicate differences with a P-value <0.05 or <0.0001, based on a two-sided t-test performed between triplicates of the indicated conditions. The maximum number of exchangeable deuterons (maxD) is indicated by a dotted line. (F) HX at 100 s of the representative peptide in the ASL of PARP2 or PARP1 with or without DNA damage is shown. *** and ns indicate differences with a P-value <0.001 or >0.05, respectively. (G) PARP1 and PARP2 structures used in panel C and D, highlighting differences between the PARP1 and PARP2 ASL during DNA engagement.

A breakthrough in the understanding of PARP1 allosteric regulation upon engaging DNA breaks came through examination of its polypeptide backbone dynamics using hydrogen/deuterium exchange coupled to mass spectrometry (HXMS)^7^. HXMS measures amide proton behavior, such that stable structures (i.e. within α-helices or the interior of β-sheets) are protected unless they undergo transient unfolding and refolding^18^. When PARP1 binds to a DNA break, three of the HD helices undergo 10000-fold faster HX^7^. This permits NAD^+^ rapid access to the catalytic center of the enzyme^19^, culminating in >1000-fold increase in PAR production^7^.

Here, we use HXMS to measure the backbone dynamics of PARP2 during DNA damage-induced activation and upon engaging different PARPi molecules. Combining this analysis with experimentation involving structure-based mutations, we uncover a distinct feature of PARP2 that is required for its full autoinhibition and generate a model for how autoinhibition is relieved when it detects DNA damage.

## Results

### PARP2 has additional DNA break-induced changes to backbone dynamics relative to PARP1

On one hand, we predicted that the homology in the allosteric HD between PARP1 and PARP2 (Fig. 1A) would lead to a similar mechanism where destabilization of HD helices accompanies activation upon binding to a DNA break. On the other hand, we recognized that the divergence throughout the shared domains, including the HD, may lead to important differences. To measure the backbone dynamic changes of PARP2 during DNA binding, we performed HXMS on PARP2 (Figs. 1B and S1A-C) in the presence or absence of a 5’ phosphorylated nicked (5’P nicked) DNA, its preferred type of DNA break^12^. We focused on the 100 s timepoint in our HXMS experiments, because it was a particularly useful timepoint in our prior PARP1 studies in reporting on protection and deprotection from HX upon engaging with a DNA break^7^. Coverage by partially overlapping peptides over the folded domains of PARP2 was complete, with gaps only in the unstructured N-terminal region (NTR). The resulting HXMS data for PARP2 were mapped onto a high-resolution cryo-EM structure of full-length PARP2 isoform 2, bound to broken DNA and HPF1 (Fig. 1C; PDB 6X0L^20^ with HPF1 removed for clarity). Binding to DNA leads to HX protection, with the strongest protection spanning the majority of the WGR (Fig. 1B,C). WGR HX protection at DNA contact points is strongest when PARP2 engages its favored substrate (5’P nicked DNA) relative to other types of breaks that we tested: an unphosphorylated nicked substrate and the favored PARP1 substrate with a gap in the DNA backbone (Figs. S1A,B and S2A). The strongest HX deprotection in PARP2 upon engaging the DNA break resides within discrete portions of the HD and one region within the ART (Fig. 1B,C). Thus, PARP2 backbone dynamics are present in all three of its folded domains (WGR, HD, and ART) upon binding to a DNA break.

When comparing with the HXMS behavior of PARP1 upon engaging with its favored substrate, a simple single-stranded break, PARP2 has clear similarities and differences (Fig. 1C,D). The WGR is similar in both PARP1 and PARP2, in that they experience strong HX protection, but the protection is more pronounced for PARP2 (Fig. 1C,D). One possibility to explain this finding is that the DNA binding surface for PARP1 is relatively more dispersed, with the WGR only representing a portion of the DNA binding surface. The WGR serves as the entire structured DNA binding surface in the case of PARP2. Additionally, PARP2 has a slower off-rate from a DNA break relative to that of PARP1^14^, correlating with the greater PARP2 WGR rigidification we observe by HXMS (Fig. 1C,D). The HD is similar in having almost the exact same positions of the αB helix and αF helix mapping with the strong HX deprotection in both PARP1 and PARP2, but PARP2 uniquely exhibits protection in much of its αE helix (Fig. 1C,D). The αE helix of PARP1 only becomes protected upon experiencing “reverse allostery” imposed by binding to substrate NAD^+19^ or a type I PARPi^15^ (Fig. S2B,C). Comparison of a nearly identical αE peptide from PARP1 and PARP2 shows that the α-helix experiences HX protection in PARP2 upon engaging a DNA break to a similar degree as in PARP1, but only after PARP1 is experiencing reverse allostery (from the PARPi, EB-47, in the comparison shown; Figs. 1E and S2B,C). In the ART of PARP2, the prominent HX deprotection upon engaging with a DNA break maps to the active site loop (ASL)(Fig. 1B,C). Deprotection of the same region in PARP1 upon binding to a DNA break is much less pronounced (Fig. 1D,F,G). Comparison of a nearly identical ASL peptide from PARP1 and PARP2 reveals that this difference is due to marked protection in the case of PARP2 when unbound to DNA in contrast to the relatively fast exchanging behavior in PARP1 (Fig. 1F). Upon binding to a DNA break, the ASL of both PARP1 and PARP2 exhibit similar HX (Fig. 1F). Our HX results are therefore consistent with PARP2 allosteric activation occurring through a combination of shared and divergent structural and dynamic changes relative to PARP1.

### HD unfolding is insufficient to permit PARP2 DNA damage-induced activation

PARP1 activation is defective when communication from the DNA binding domains is broken and HD unfolding is abolished (i.e. in the W318R mutation in the F3 domain of PARP1^7^). Lacking domains F1-F3, the only structured DNA binding domain of PARP2 is the WGR. Thus, we predicted that interrupting communication from the WGR would abolish HD unfolding in a PARP2 mutant. To test this by HXMS, we first needed to identify such a PARP2 mutation. Of the several WGR mutations in PARP2 that are known to eliminate DNA damage-induced activation, only one that we are aware of, N116A, retains high affinity for a DNA break^12,13,21^. N116 is located in WGR β2 strand that approaches the HD at the C-terminal portion of αE helix and a loop that is C-terminal to it^12,13,16^. The conserved WGR residue in PARP1 (N567) interfaces with F1 and the HD and abolishes catalysis when mutated to an alanine^22^, and was predicted to have the same impact on PARP2^12^. In designing the experiment, we considered our finding that WT PARP2 has nearly complete HX by 100 s (Figs. 1B and S1A,B) in the regions we sought to examine. Thus, we instead chose an earlier timepoint (10 s) for increased sensitivity in comparing WT to N116A (Figs. 2 and S3). As predicted, the N116A mutation has almost identical HX protection at the portions of the WGR that directly contact broken DNA (Figs. 2A and S3C,D), but portions of the WGR that connect from N116 in WT PARP2 to the HD (and HD αE helix [Fig. 2D]) exhibit reduced HX protection (Figs. 2A and S3B). Contrary to our prediction, the HD helices that are strongly destabilized in WT PARP2 (measured by strong deprotection from HX) upon binding to a DNA break are destabilized to a similar extent in N116A (Fig. 2A-C,F,G). In contrast to the modest impact on HD destabilization, we noted that the destabilization of the ASL Helix within the ART was nearly abolished in the N116A mutant (Fig. 2A,D-G). The inability of the N116A mutant to undergo DNA damage-induced enzymatic activation^12^, together with extensive HD destabilization (Fig. 2), indicate that unlike PARP1, HD destabilization is insufficient to relieve autoinhibition. The inability of the N116A mutant to undergo DNA damage-induced enzymatic activation^12^ together with a lack of ASL Helix destabilization and reduced HD destabilization (Fig. 2A,C,F) indicate that destabilization of the ASL Helix is required to destabilize the HD to the degree of DNA-bound WT PARP2. Unlike in PARP1^7^, HD destabilization is insufficient to relieve PARP2 autoinhibition. Rather, our HX experiments indicate the likelihood that a second destabilization event, namely destabilization of the ASL Helix, is needed to relieve autoinhibition.

**Figure 2:**
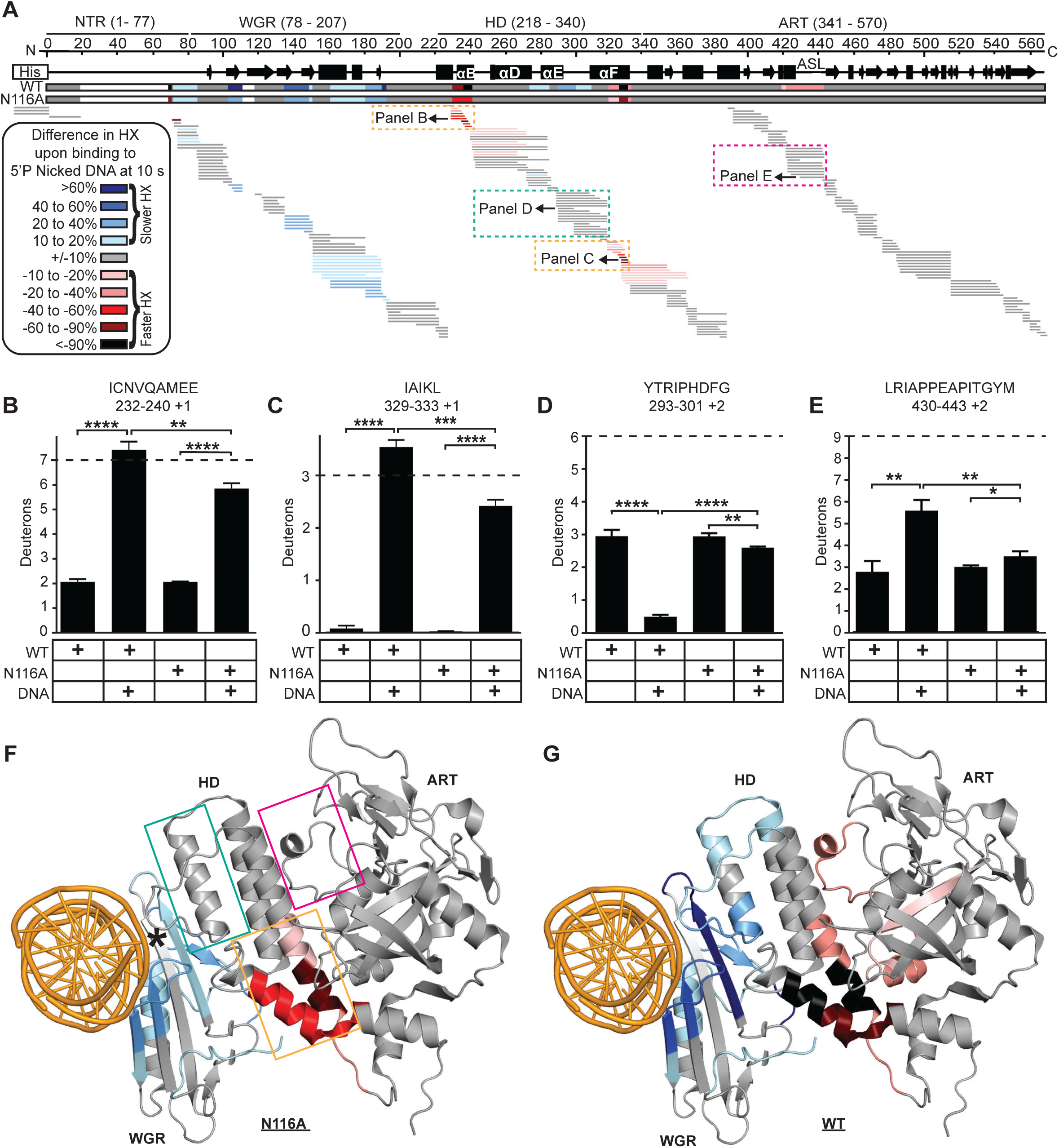
Extended contacts between the WGR β-sheet and the HD confer ASL Helix destabilization upon PARP2 engaging a DNA break. (A) HX difference plot between N116A PARP2 with and without 5’P nicked DNA at 10 s. The consensus behavior at each N116A PARP2 residue is displayed in a horizontal bar below the secondary structure annotation, along with a horizontal bar representing WT PARP2 data for comparison. (B-E) HX at 10 s of the indicated representative peptides in the HD αB helix (B), HD C-terminus of αF helix (C), HD C-terminus of the αE helix (D), or ASL (E) for the indicated conditions. *, **, ***, and **** indicate differences with a P-value <0.05, <0.01 <0.001 or <0.0001, respectively, based on a two-sided t-test performed between triplicate samples of the indicated conditions. (F) Consensus HX data from N116A PARP2 with or without 5’P nicked DNA in panel A mapped on the cryo-EM structure of PARP2 complexed with DNA and HPF1 (PDB 6X0L; displayed as shown in Fig. 1C). The position of N116 is indicated with an *. (G) Consensus HX data from WT PARP2 upon binding to 5’P nicked DNA in panel A mapped on the cryo-EM structure of PARP2/DNA/HPF1 complex (PDB 6X0L; displayed as shown in Fig. 1C).

### HD contacts stabilize the ASL Helix to autoinhibit PARP2

We examined high-resolution structural data to assess the shape of the ASL in the presence or absence of close HD contacts. Several structural models of PARP2 include the HD in close proximity to the ASL region^20,23–26^. In these structures, the ASL has a backbone geometry that is readily assigned as a short α-helix (denoted here as ASL Helix in Fig. 3A) by Pymol. Other structural models exist of the ART in isolation^7,27^ or where the HD is present but separated due to an extended overall PARP2 configuration^21^. In these structures, the absence of the HD is accompanied by an ASL with a backbone path that lacks clear α-helical geometry (Fig. 3A). Thus, existing structural models support the idea that PARP2 has a stable α-helix at its ASL region that requires close association of the HD to maintain its rigid secondary structure.

**Figure 3:**
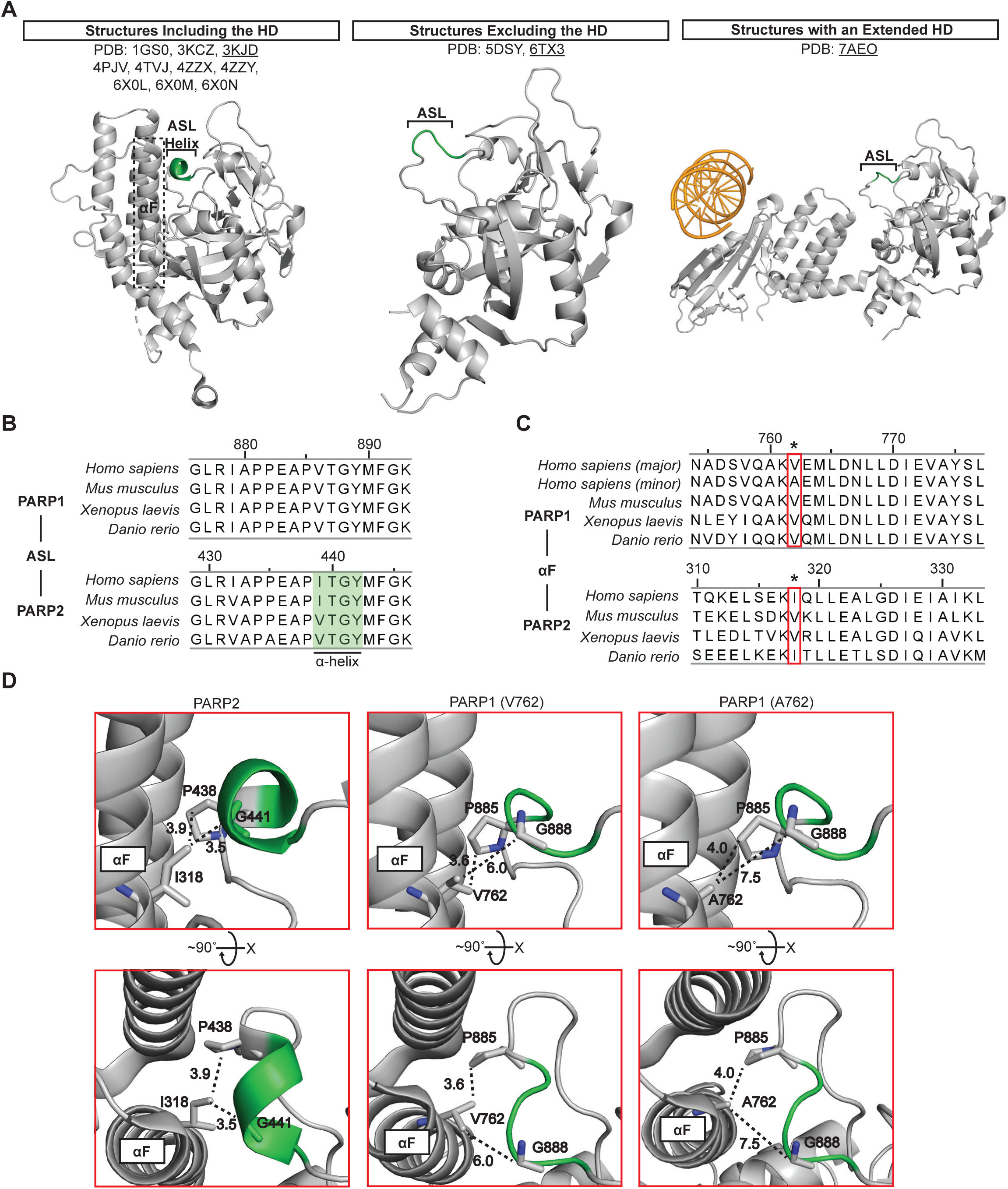
The PARP2 active site loop forms an α-helix upon stabilization through contacting the HD. (A) The 3D structures with the HD (αF helix) contacting the ART, excluding the HD, or with the HD distant from the ASL are shown. (B-C) Sequence alignment of ASL with α-helical residues overlayed in green (B) or sequence alignment of HD αF (C) of PARP1 and PARP2 in humans and in selected orthologs. (D) Crystal structures in corresponding orientations highlighting the unique contacts between ASL and αF of PARP2 (PDB 3KJD) and PARP1 (PDB 3GN7) are shown. V762 PARP1 model (middle) was visualized using PDB 3GN7 and the Pymol mutagenesis wizard.

The ASL Helix in PARP2 is unlikely to be due to any sequence changes. There is a single residue substitution between human PARP1 and PARP2 (corresponding to V886 and I439, respectively) in the residues that form the short α-helix or are located nearby (Fig. 3B). Stability from the HD in PARP2 likely comes from the packing of the ASL region with the HD, especially the αF helix (Fig. 3A,C). We were particularly intrigued by the recent surprising report that a mutation, I318A, in the αF helix of PARP2 apparently reduces autoinhibition^16^. This PARP2 residue corresponds to PARP1 position 762 that has two substantially represented variations in the human population: a valine as the major form, and an alanine as the minor form^28,29^ (Fig. 3C,D). In the absence of DNA, A762 PARP1 is strongly autoinhibited^7^. Thus, an alanine at the same position has no measurable impact on PARP1 but leads to a clear defect in autoinhibition for PARP2. In PARP1, either V762 or A762 in the two major SNP forms only contacts P885 at the N-terminal end of the ASL, with the C-terminal end of the ASL extending too far away (Fig. 3D). I318 is clearly engaged with the ASL Helix in PARP2, with the I318 side-chain abutting P438 at the N-terminal end of the ASL and the α-carbon of G441 at the C-terminal end (Fig. 3D). The I318A mutation, therefore, likely reduces the hydrophobic contacts with both ends of the ASL Helix. Thus, we predicted that I318A would have reduced ASL Helix stability, relative to WT PARP2.

To test this prediction, we performed HXMS on the I318A mutant protein and compared it to the HX measured for WT PARP2 (Fig. 4A). The only region with any substantial change was a discrete region of the ART that spans the ASL Helix, where the I318A mutant exhibited faster HX (Fig. 4A-C). Thus, the ASL is destabilized by the I318A mutation, helping to explain its reported defect on autoinhibition^16^. Notably, no corresponding destabilization of HD helices accompanied the I318A mutation (Fig. 4A-C)(as it does for WT PARP2 upon engaging a DNA break [Fig. 1]), indicating that destabilization of the ASL Helix can occur without HD destabilization. We previously found that the so-called ΔHD mutation of PARP2 that replaces the αD, αE, and αF helices with a flexible linker abolishes autoinhibition^7^. The ΔHD mutation, thus removes any steric blocking of NAD^+^ access^7,19^ as well as removing the ASL contact with the αF helix that includes I318. Using an assay for HPF1-dependent serine automodification, we find that WT PARP2 is robustly autoinhibited in the absence of a DNA break and I318A PARP2 has a measurable defect in autoinhibition that can be seen with activity at later timepoints (Fig. 4D). In contrast, ΔHD PARP2 exhibits robust activity from the earliest timepoints (Fig. 4D). Together, these findings indicate that ASL destabilization in PARP2 alone is sufficient to drive some leaky poly(ADP)- ribosylation activity in the absence of DNA, but that complete relief from autoinhibition requires the additional step of HD unfolding.

**Figure 4:**
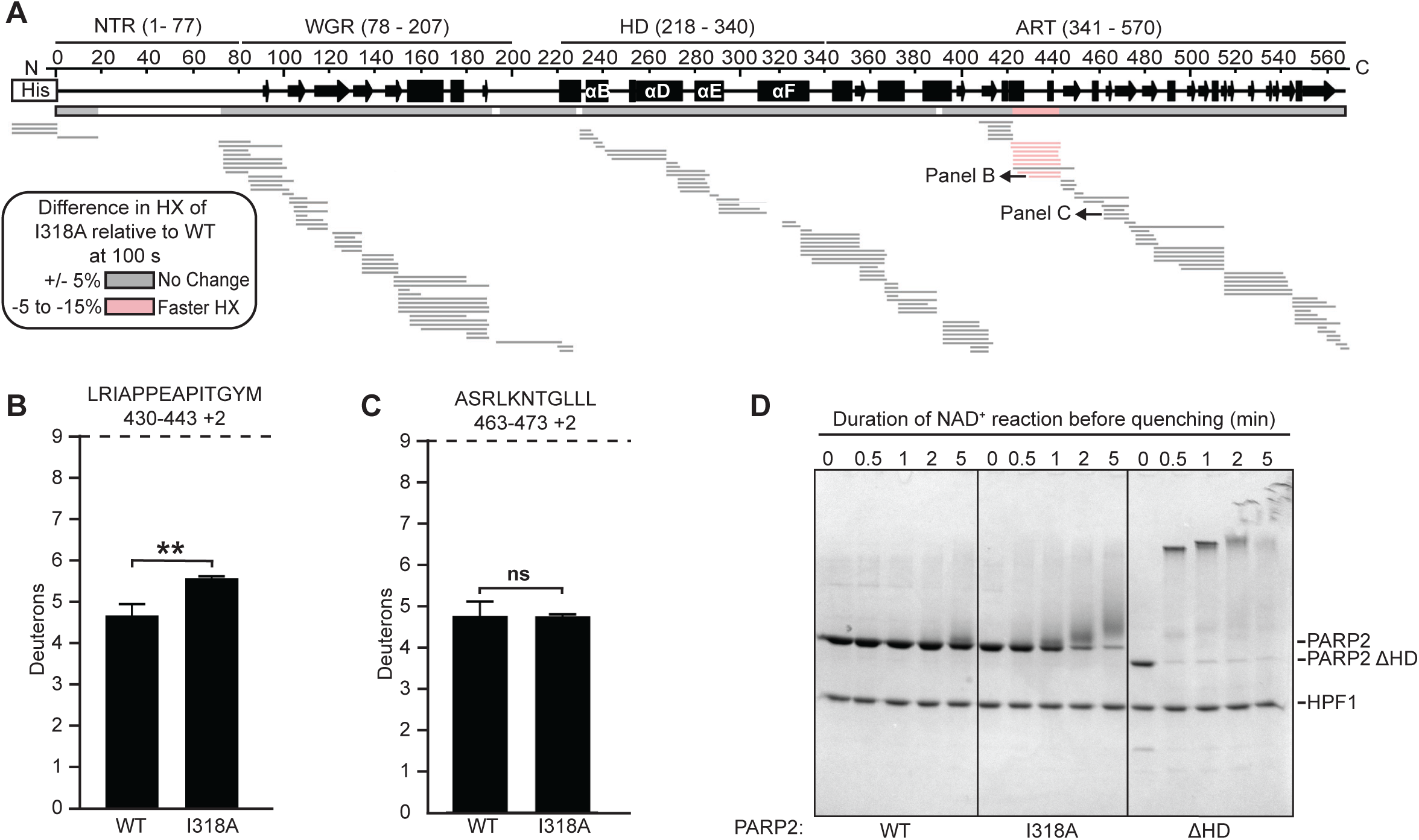
Contacts between the HD and the ASL Helix maintain PARP2 autoinhibition until their separation via DNA damage engagement. (A) HX difference plots between WT PARP2 and I318A at 100 s. Note that both WT and I318A are measured in the absence of DNA. (B) HX at 100 s of the representative peptide of ASL in WT PARP2 and I318A. (C) HX at 100 s of a representative peptide in an unchanging region of the ART in WT PARP2 and I318A. (D) Catalytic activity of WT PARP2, I318A PARP2, and ΔHD PARP2 (1 μM) in the presence of NAD^+^ (500 μM) was measured using an SDS-PAGE automodification assay.

### Olaparib and EB-47 stabilize the ASL Helix while inhibiting PARP2

Since all FDA-approved clinical PARPi target the NAD^+^-binding site at the catalytic center of both PARP1 and PARP2, and since this is juxtaposed to the ASL Helix location, we reasoned that some PARPi may stabilize the ASL Helix. We tested this notion with HXMS and using PARP2 that is engaged with a DNA break and one of each of the PARPi molecules (Figs. 5, S4 and S5). Most PARPi molecules did not confer a change in HX to the ASL Helix, but Olaparib drove measurable protection to the ASL Helix (Figs. 5, S4, and S5). EB-47, a non-clinical PARPi that mimics NAD^+^-binding also conferred some HX protection to the ASL Helix (Figs. 5C and S4A). In the allosteric HD domain and throughout the DNA-binding WGR of PARP2, we only observed very small regions of HX protection that included the WGR β1 sheet upon binding to Talazoparib, Niraparib, Rucaparib, UKTT15, and EB-47, and mild HX deprotection upon binding to Olaparib (Fig. 5). The HX behavior of the HD of PARP2 with various PARPi (Figs. 5 and S4) contrasts with that of PARP1^15^. This behavior is also consistent with the emerging view that the impact of PARPi on PARP2 and its DNA binding behavior is largely driven by subtle structural changes^16^, rather than like in PARP1 where the backbone dynamics of the HD and DNA binding domains is heavily influenced by the particular type of PARPi^15^. In all, though, the most notable change in PARP2 that varies by the particular PARPi molecule is the ASL Helix. We envision that this occurs due to either direct binding of Olaparib or EB-47 to the ASL or restricting movement of the αF helix within the HD that, in turn, leads to greater stability of the ASL Helix through its own direct contacts (Fig. 3). It will be interesting in the future to investigate whether or not the ASL Helix could provide a ‘handle’ that is only available in PARP2 and not PARP1 for identification or design of new PARPi molecules that are specific for PARP2 functions^30,31^.

**Figure 5:**
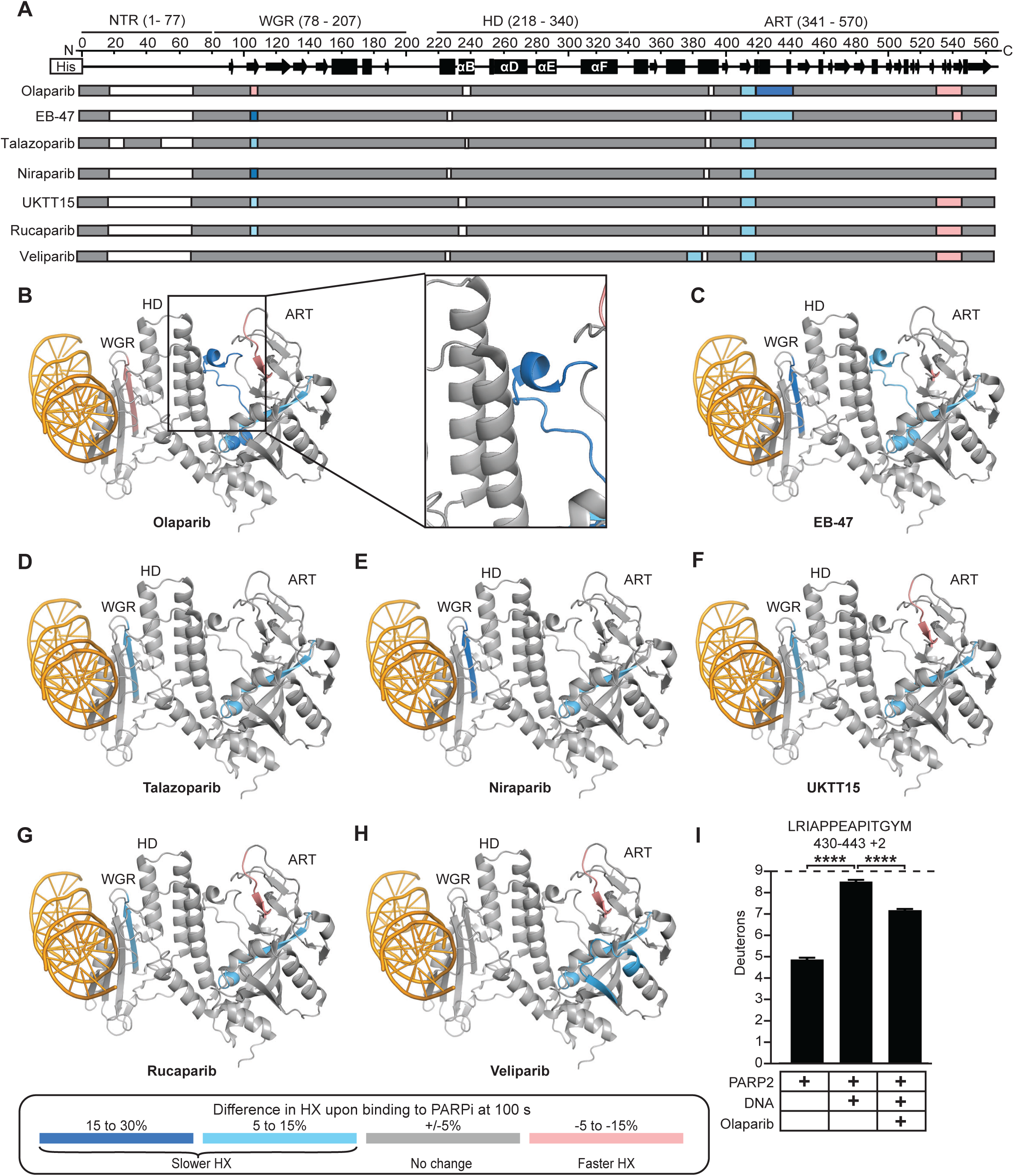
Olaparib and EB-47 stabilize the ASL of PARP2. (A) Consensus HX between PARP2 with 5’P nicked DNA and PARP2 complexed with 5’P nicked DNA and the indicated PARPi at 100 s. (B-H) Consensus HX data from Panel A mapped on the cryo-EM structure of PARP2/DNA/HPF1 (PDB 6X0L; displayed as shown in Fig. 1C). (I) HX at 100 s of the representative ASL peptide of PARP2 with the indicated conditions is shown. **** indicates a difference with a P-value <0.0001, based on a two-sided t-test performed between triplicate samples of the indicated conditions.

## Discussion

Using HXMS as a primary experimental approach, our study reveals that the autoinhibition of PARP2, and relieving it upon DNA break-induced enzymatic activation, has additional allosteric mechanisms that were not predictable based on existing structural models or comparisons to PARP1. By analogy with an automobile with two axles (Fig. 6A; “PARP car”), PARP1 has a brake on one axle while PARP2 has brakes on both. Here, we found with PARP2 that there is indeed communication from the WGR upon binding a DNA break, that, as in PARP1, leads to massive destabilization of multiple HD α-helices, including the αF helix that would otherwise sterically block access of substrate, NAD^+^. Unlike in PARP1, though, in PARP2 the WGR is strongly rigidified throughout most of the domain including its uniquely extended β-sheet that abuts the HD αE helix (Fig. 6B). This generates rigidity to one portion of the HD (αE helix) simultaneously with the flexibility imparted to another portion (especially the αB helix and the C-terminal half of the αF helix). It is reasonable to conclude that HD rigidity to the αE helix accompanies movement of the HD away from the ART^16,20,21^ that, in turn, removes contacts that stabilize the ASL Helix. We found that the ASL Helix, unique to PARP2, must remain intact to restrict leaky enzymatic activity. Reciprocally, we found using a mutant (N116A) that does not permit ASL Helix destabilization, that is able to silence PARP2 even when there is robust destabilization of the HD helices, including the αF helix. The ASL Helix is directly N-terminal to a residue (M443) contacting NAD^+^ involved in catalysis^32^, and likely influences the position of other neighboring NAD^+^ binding residues directly adjacent to it^32,33^. Thus, we envision that it must at least transiently sample unfolded states for productive NAD^+^ binding during catalysis. Our findings emphasize that even the closest relative of PARP1 in the PARP superfamily of enzymes can have unique mechanisms critical to their tight autoinhibition and massive activation to achieve their function in signaling a DNA break.

**Figure 6:**
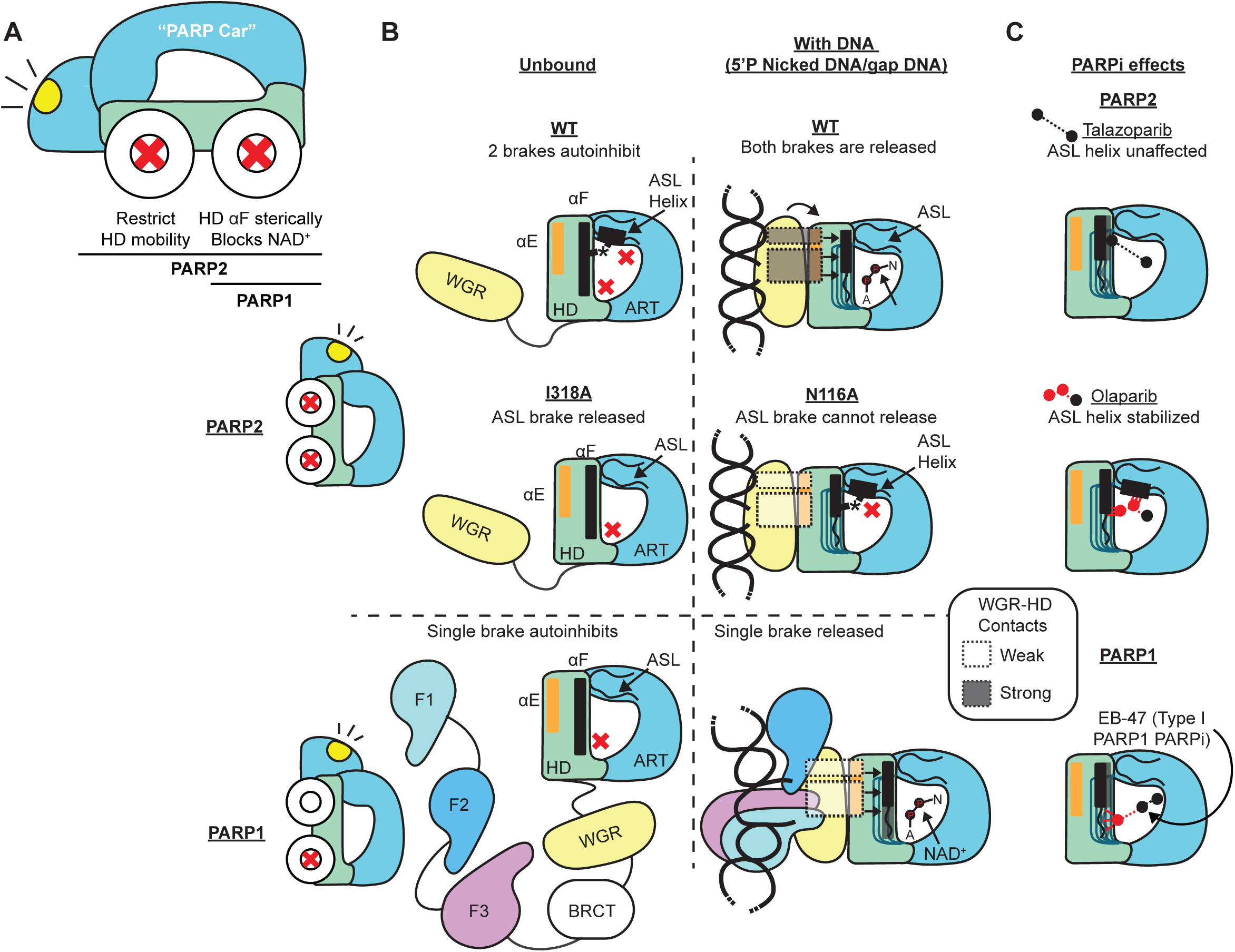
PARP2 undergoes an additional forward allosteric pathway during DNA dependent activation that is distinguished from PARP1. (A) Analogy of PARP1 and PARP2 to a “PARP car”. Models of PARP1 and PARP2 comparing autoinhibition (B) and PARPi response within the CAT (C).

Our findings with HXMS complement what has been learned by indispensable structural models achieved by crystallography and cryo-EM^20,21^, and extend the understanding further about PARP2 activation. For the unfolding of the HD, this was predicted based on the finding that removing the key α-helices for PARP1 activation leads to constitutive activity even in the absence of a DNA break for both PARP1 and PARP2^7^. This notion was further supported by minor states in single particle cryo-EM analysis of PARP2 engaged with DNA ends, where structural models were only at moderate resolution (6-7 Å), but nonetheless demonstrated populations of PARP2 molecules where the HD moved away from the WGR^20^. Likewise, a more extreme extension was captured in a crystal form of PARP2 bound to a DNA break where the key HD helices were dramatically reconfigured (PDB 7AEO^21^). The rapid HX we observe in solution of the key HD helices upon PARP2 activation, supports a model where these helices are rapidly sampling unfolded states since the rapid HX observed by 10 s (Fig. 2) happens as a consequence of the α-helices unfolding and refolding several times in that short timeframe. Every time they unfold, the block to NAD^+^ access created by the αF helix is removed. By analogy to PARP1^34^, the increased HD flexibility in PARP2 is likely coupled to transient structural motions/reconfigurations nearby. The prior studies did not, however, anticipate the role that we report of the ASL Helix in PARP2 autoinhibition, and how its unfolding is required even if HD flexibility occurs (Fig. 2). Further, the leaky activity of the I318A (Fig. 4D) in the absence of detectable HX deprotection of the αF helix (Fig. 4A), which provides the primary steric block to NAD^+^ binding^7^, suggests that PARP2 permits a slightly higher amount of baseline NAD^+^ binding relative to PARP1. We envision that the rigid ASL Helix in PARP2 is used to restrict structural mobility of the HD. When the HD is released by DNA binding and ASL Helix melting, then NAD^+^ binding becomes fully productive for catalysis (Fig. 6).

There has been a strong indication of redundancy in PARP1 and PARP2, which our study supports (including the near perfect match in the boundaries of HD unfolding upon binding to a DNA break; Fig. 1). On the other hand, our study strongly adds to the growing list of evidence to indicate they have important independent functions that need to be considered, especially when targeting them with small molecule inhibitors in the clinic. The fact that removal of both is required to produce a lethal phenotype in mice^11^, as well as the findings that they have similar, but not identical, abilities to rapidly bind and be activated by DNA breaks, argue for redundancy. Single knockouts of PARP1 and PARP2 lead to high sensitivity to genotoxic treatments^11,35^ indicating that they cannot sufficiently compensate for each other when challenged. We speculate that the different aspects of DNA binding, autoinhibition, and activation by DNA breaks by PARP1 and PARP2 are related to the non-overlapping roles they play in DNA damage detection and signaling.

The issues of redundancy versus independent functions of PARP1 and PARP2 is a pressing issue in light of findings emerging from the clinical use of PARPi. The current FDA-approved PARPi are well tolerated as single agents, but toxicity to healthy tissues, especially the bone marrow, becomes unmanageable when combined with therapies that target other pathways^36–38^. The combination therapies could otherwise be very helpful because they, in principle, lead to more potent tumor cell killing and delay or diminish PARPi resistance emerging in patients undergoing treatment^37^. One hopeful solution is with PARPi compounds that are selective for PARP1^39,40^, with the underlying premise that the bone marrow toxicity with the first generation of PARPi is due to disrupting both PARP1 and PARP2 function in healthy cells. PARP2-specific PARPi have garnered less attention^30^ but our study would add to the motivation to pursue them (Fig. 6C). They may have similar benefits as PARP1-specific PARPi, and they may also have specific utility in some clinical contexts. Indeed, a benefit of PARP2-specific PARPi has been proposed in B-cell lymphoma^31^ and prostate cancer^30^. Structural studies, including solution-based measurements like one can achieve with HXMS, will surely continue to be useful in assessing strategies to achieve desired outcomes with both PARP1-specific and PARP2- specific small molecule interventions. HXMS is sensitive enough to detect even very subtle changes conferred by different PARPi molecules^41^. In addition, the structural framework and methodology we present on PARP2 serve as a useful starting point to design and experimentally assess new PARPi molecules into the future.

### Experimental Procedures

#### Expression Constructs and Mutagenesis

FL PARP2 isoform 2 (1-570) was expressed from a pET28 vector with an N-terminal hexahistidine tag^9,42^. FL human HPF1 was expressed from the pET28 vector with an N-terminal SMT sumo-like His-tag^9^. For I318A and N116A PARP2, site-directed mutagenesis was performed on the FL PARP2 isoform 2 (1-570) in the pET28 vector using the QuikChange protocol (Stratagene) and verified by Sanger sequencing. PARP2 ΔHD was generated by deleting residues 260-334 from the pET28 vector containing FL PARP2 isoform 2 (1-570)^7^.

#### Protein Expression and Purification

WT, I318A, and N116A PARP2, HPF1, and WT PARP1 were expressed in Rosetta 2 (DE3) *E. coli* cells in media supplemented with 10 mM benzamide. Ni^2+^ affinity, heparin, and gel filtration chromatography were used to purify WT, I318A, and N116A PARP2 and WT PARP1, as described^42^. HPF1 was purified as described^9^.

#### Hydrogen/Deuterium Exchange Mass Spectrometry (HXMS)

Prior to deuterium on-exchange at room temperature (RT), 6 μM of PARP2 was incubated for 30 min with or without 12 μM of dumbbell DNA with a single nucleotide gap (5’-GCTGAGCTTCTGGTGAAGCTCAGCTCGCGGCAGCTGGTGCTGCCGCG-3’) or 12 μM of dumbbell nicked DNA with a 5’OH (5’- GCTGAGCTTCTGGTGAAGCTCAGCTCGCGGCAGCTGGTGCTGCCGCGA-3’) or 12 μM of dumbbell DNA with a 5’P nick (5’P- GCTGAGCTTCTGGTGAAGCTCAGCTCGCGGCAGCTGGTGCTGCCGCGA-3’). For the PARP2+DNA±PARPi experiments, 12 μM of EB-47, Talazoparib, Niraparib, Olaparib, Rucaparib, Veliparib, or UKTT15 was added to the PARP2 + 5’P nicked DNA mixture and incubated for another 30 min. For each experiment, deuterium on-exchange was performed at RT in triplicate (n=3) by adding 5 μL of protein I318A, N116A, or WT PARP2 to 15 μL of deuterium on-exchange buffer (10 mM HEPES, pD 7.0, 100 mM NaCl, in D_2_O, pD = pH + 0.4138) to yield a final D_2_O concentration of 75%. After 10 or 100 s, the exchange reaction was quenched by mixing the deuterated sample (20 µL) into 30 μL of ice-cold quench buffer (1.66 M guanidine hydrochloride, 10% glycerol, and 0.8% formic acid to make a final pH of 2.4-2.5) and immediately frozen in liquid nitrogen until further use. The non-deuterated (ND) samples of PARP2 were prepared in 10 mM HEPES, pH 7.0, 100 mM NaCl buffer and each 20 μL aliquot was quenched into 30 μL of quench buffer. To mimic the on-exchange experiment, the fully deuterated (FD) samples were prepared in 75% D_2_O and denatured under acidic conditions (0.5% formic acid). FD Samples were incubated for 48 hours to ensure every amide proton along the entire polypeptide chain could undergo full exchange, and a 20 µL aliquot is quenched with 30 µL ice-cold quench buffer (1.66 M guanidine hydrochloride, 10% glycerol, and 0.8% formic acid to make a final pH of 2.4-2.5). 50 μL samples were thawed at 0°C, rapidly injected into pepsin column, and pumped at initial flow rate of 50 μL min^-1^ for 2 min followed by 150 μL min^-1^ for another 2 min. Pepsin (Sigma) was coupled to POROS 20 AL support (Applied Biosystems) and the immobilized pepsin was packed into a 64 μL column (2 mm × 2 cm, Upchurch). The pepsin-digested peptides were trapped onto a TARGA C8 5 μm Piccolo HPLC column (1.0 × 5.0 mm, Higgins Analytical) and eluted through an analytical C18 HPLC column (0.3 × 75 mm, Agilent) with a 12-100% buffer B gradient at 6 μL/min (Buffer A: 0.1% formic acid; Buffer B: 0.1% formic acid, 99.9% acetonitrile). The effluent was electrosprayed into the Q-Exactive (Thermo Fisher Scientific) and the MS data acquisition over the mass range 200-2000 m/z were acquired at 70,000 resolution. The effluent was electrosprayed with ion spray voltage of 3.5 kV and capillary temperature operated at 250 °C.

#### PARP2 Peptide Identification

For peptide identification the ND samples were injected into the Q- Exactive (Thermo Fisher Scientific) for tandem mass spectroscopy (MS/MS). Scan range for MS/MS data was 200-2000 m/z at 17,500 resolution, where the ions were fragmented by CID with normalized collision energy 30-35. The raw MS/MS files were processed with Proteome Discoverer v 2.4 (Thermo Fisher Scientific). Database search was performed using SEQUEST with a peptide tolerance of 8 ppm and a fragment tolerance of 0.1 Da against an extensive sequence database containing the sequence of PARP2, pepsin and other in-house contaminants identified in prior HXMS studies. We specified pepsin as the protease in our search as a non-specific digestion enzyme. An FDR of 10% was employed for the protein identification. After MS/MS was performed for the first ND sample, the sequence of the peptide, RT, charge, m/z, confidence etc., were used to generate an exclusion list. While performing MS/MS on the second ND sample, the instrument employed this exclusion list to collect the MS2 scan of the less intense peptides that were not identified in the previous ND sample. At least 2 such exclusion lists were generated to increase the number of unique peptides and sequence coverage of the protein. Later, peptide information based on the above MS/MS of all NDs were merged to have a final peptide pool.

#### HXMS Analysis and Plotting

HDExaminer software (v 2.5.0) was used, which uses the peptide pool information to identify the deuterated peptides for every sample in the HXMS experiment. The quality of each peptide was further assessed by manually checking mass spectra. The level of HX of each reported deuterated peptide is corrected for loss of deuterium label (back-exchange after quench) during HXMS data collection by normalizing to the maximal deuteration level of that peptide in the FD samples. After normalizing, we then compared the extent of deuteration to the theoretical maximal deuteration (maxD, i.e. if not back-exchange occurs). The median extent of back-exchange in our experiments was 29% (Fig. S1C). The data analysis statistics for all the protein states are in Supplementary Data. The difference plots for the deuteration levels between any two samples was obtained through an in-house script written in MATLAB. The script compares the deuteration levels between two samples (e.g. PARP2 and PARP2 with 5’P nicked DNA) and plots the percent difference of each peptide, by subtracting the percent deuteration of PARP2 with 5’P nicked DNA from PARP2 and plotting according to the color legend in stepwise increments. The plot of representative peptide data is shown as the mean of three independent measurements +/- SD. Statistical analysis included a t-test with a P-value <0.05. HX experiments of PARP1 with or without DNA^7^ and/or EB-47^15^ have been published. To compare PARP1 and PARP2 datasets, HX samples of PARP1 were repeated in triplicate to have the same peptide digestions and subsequent peptide data, and HX changes in HD peptides were compared between PARP1 and PARP2 with the indicated conditions. HXMS data at 100 s for PARP2 and in the presence of gap DNA, 5’OH nicked DNA, and 5’P nicked DNA was plotted through an in-house script written in R (Fig. S1A).

#### SDS-PAGE assay

Auto-modification of PARP2 was performed using 1 μM PARP2 (WT/I318A/ΔHD), 1 μM HPF1, and 500 μM NAD^+^ for various time points. Prior to resolution on an 8% or 12% SDS-PAGE, each reaction was quenched by adding SDS-PAGE loading buffer. Gels were visualized using Imperial stain.

## Acknowledgements

We thank our UPenn colleagues N. Bryan and L. Mayne for discussions on HXMS experiments. This work was supported by NIH grant CA259037 (to B.E.B., J.M.P., and T.T.T.) and an Early Career Award from Basser Center for BRCA (to R.B.).

## Author contributions

E.S.S., R.B., J.M.P., and B.E.B conceived the study. E.S.S., R.B., M-.F.L., J.M.P., and B.E.B. designed the experiments and analyzed the data. E.S.S., R.B., and M-.F.L. performed the experiments. E.S.S., R.B. T.T.T., and M-.F.L. provided key reagents. E.S.S. and B.E.B. wrote the manuscript. All authors edited the manuscript.

## Data and code availability

The HXMS data in this study has been deposited to the PRIDE database (accession code: PXD039406). Custom code used in this study has been deposited in Dryad (doi:10.5061/dryad.qjq2bvqq6).

## Supplementary figure legends

**Figure S1 (Related to Fig. 1):**
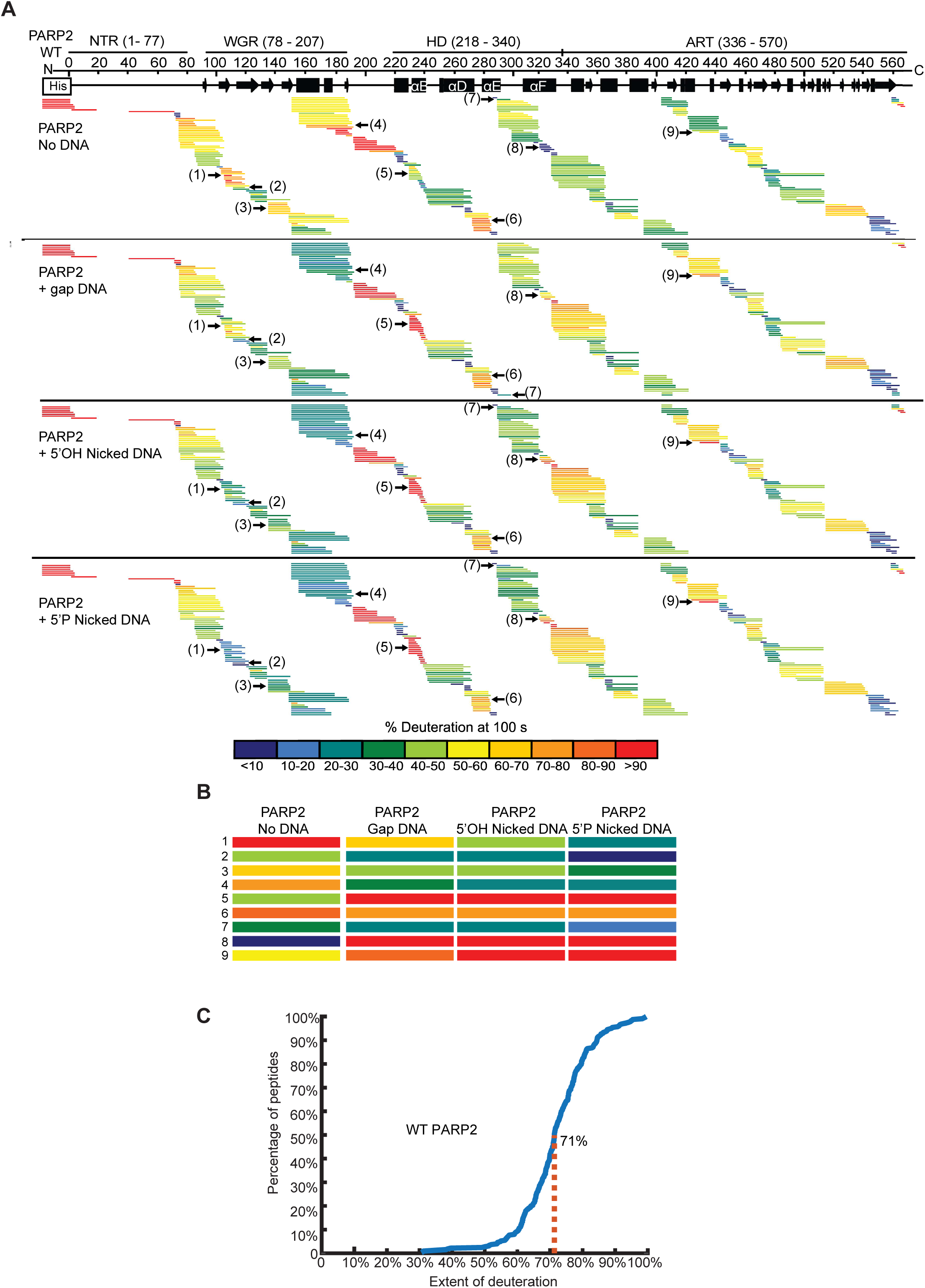
Ribbon plots for HX rates of PARP2 with or without indicated DNA damage types at 100 s and extent of deuteration in a FD control sample. (A) HXMS data at 100 s for PARP2 with or without gap DNA, 5’OH nicked DNA, or 5’P nicked DNA. Gaps represent the missing coverage. The HX is shown as % deuteration. (B) Peptides 1-9 in panel A are enlarged and compared to highlight key differences in HX between indicated conditions at regions when PARP2 binds to different DNA damages. (C) Distribution curve of a representative FD sample is shown.

**Figure S2 (Related to Fig. 1):**
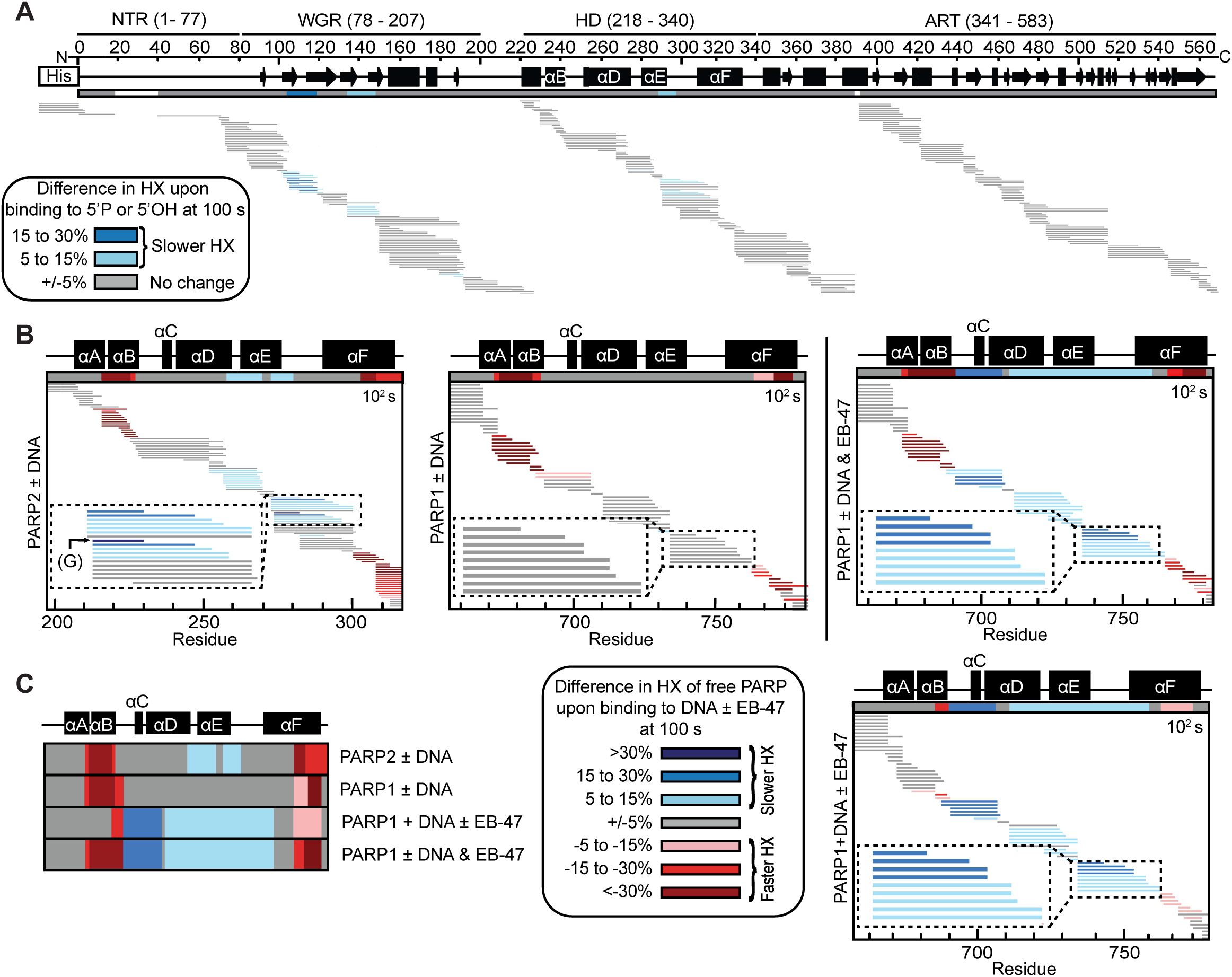
HX changes for WT PARP2 upon binding to 5’P nicked DNA or 5’OH nicked DNA and PARP1 or PARP2 HD upon binding to DNA and/or EB-47. (A) HXMS difference plots between PARP2 with 5’P nicked DNA or 5’OH nicked DNA at 100 s. Horizontal bars represent PARP2 peptides. White gaps represent the missing coverage. (B) Percent difference in HX upon binding DNA, EB-47, or both (as indicated) in the HD region for PARP2 and PARP1 at 100 s. (C) Consensus HX behavior from panel B is displayed.

**Figure S3 (Related to Fig. 2):**
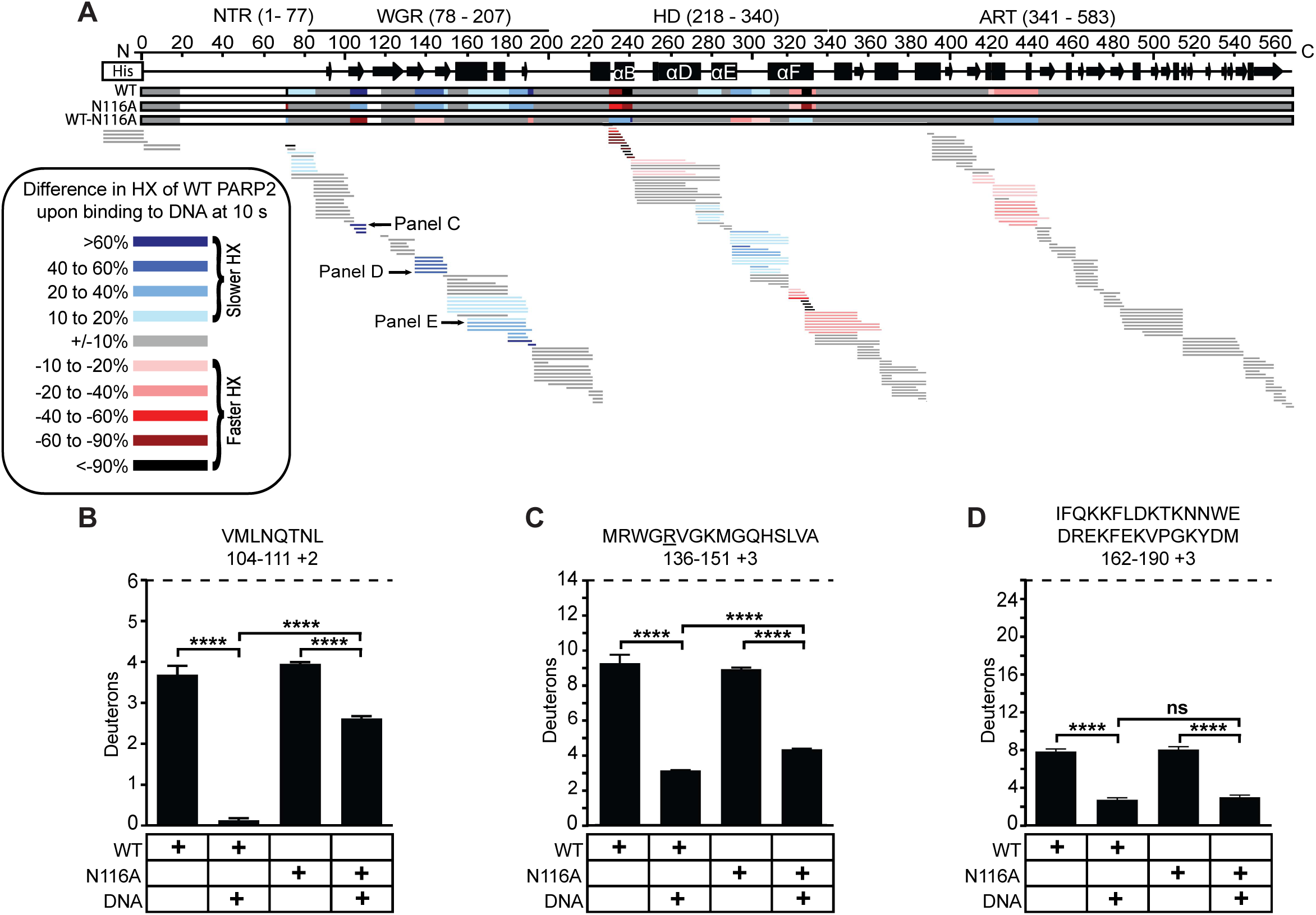
HX changes for WT or N116A PARP2 upon binding to 5’P nicked DNA. (A) HXMS difference plots between WT PARP2 and WT PARP2 with 5’P nicked DNA at 10 s. The consensus behavior at each WT PARP2 residue is displayed in a horizontal bar below the secondary structure annotation, along with a horizontal bar representing PARP2 N116A data for comparison. The difference in HX between WT PARP2 with 5’P nicked DNA and N116A PARP2 with 5’P nicked DNA is displayed in the third horizontal bar. (B-D) HX at 10 s of the representative WGR β2 strand peptide (B), WGR β3 strand peptide, with R140 underlined (C), and representative WGR α-helical DNA binding residues peptide (D), of WT or N116A PARP2 with or without 5’P nicked DNA, is shown. **, ***, and **** indicate differences with a P-value <0.01, <0.001 or <0.0001, respectively, while ns means the difference in values is not significant, based on a two-sided t-test performed between triplicate samples of WT or N116A PARP2 with or without 5’P nicked DNA.

**Figure S4 (Related to Fig. 5):**
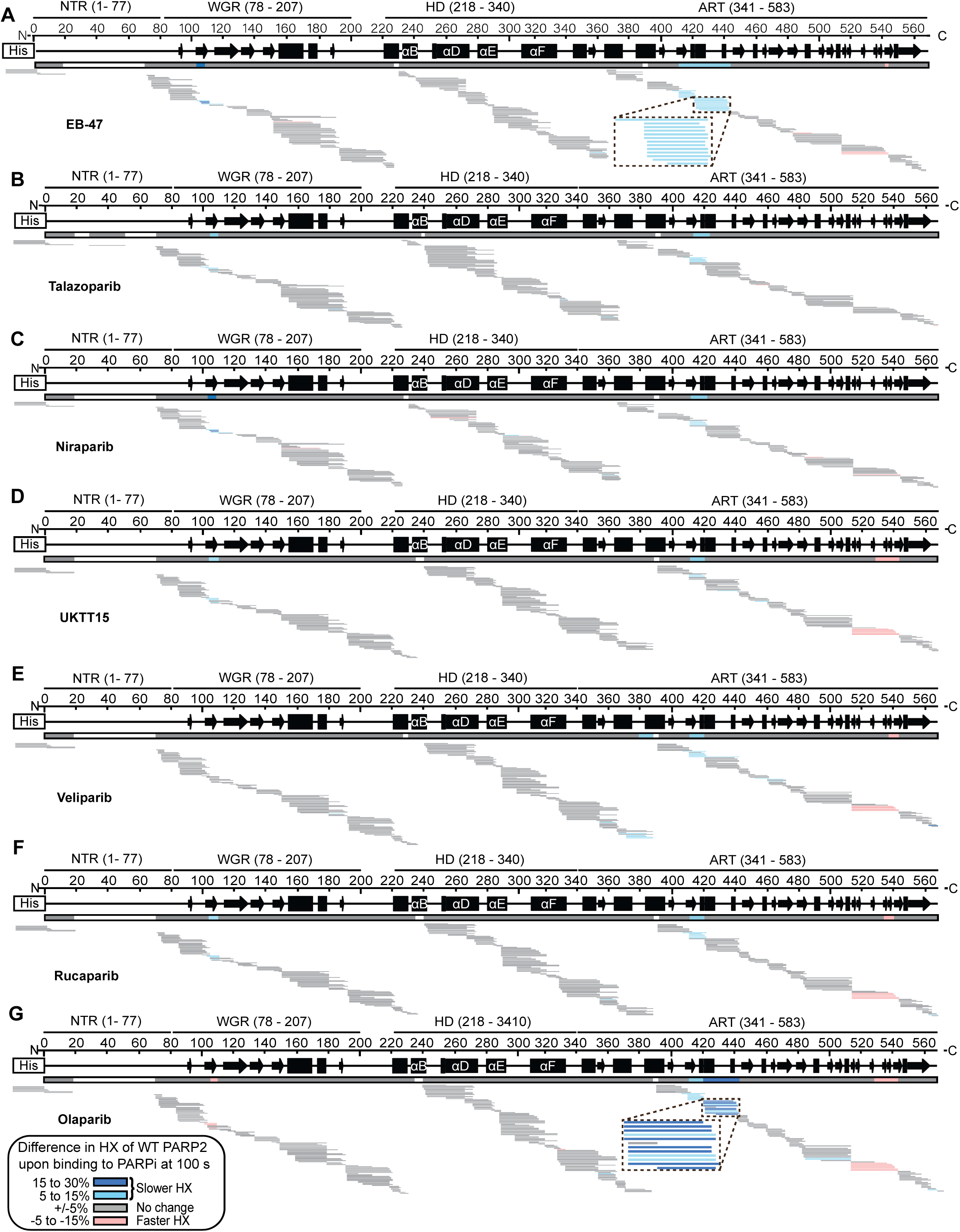
Difference plots representing the HX changes for WT PARP2 complexed with 5’P nicked DNA upon binding to PARPi. (A-F) HX difference plot between WT PARP2 with 5’P nicked DNA and the indicated PARPi at 100 s. Horizontal bars represent PARP2 peptides. White gaps represent the missing coverage. The color key of HX differences is shown below panel G.

**Figure S5 (Related to Fig. 5):**
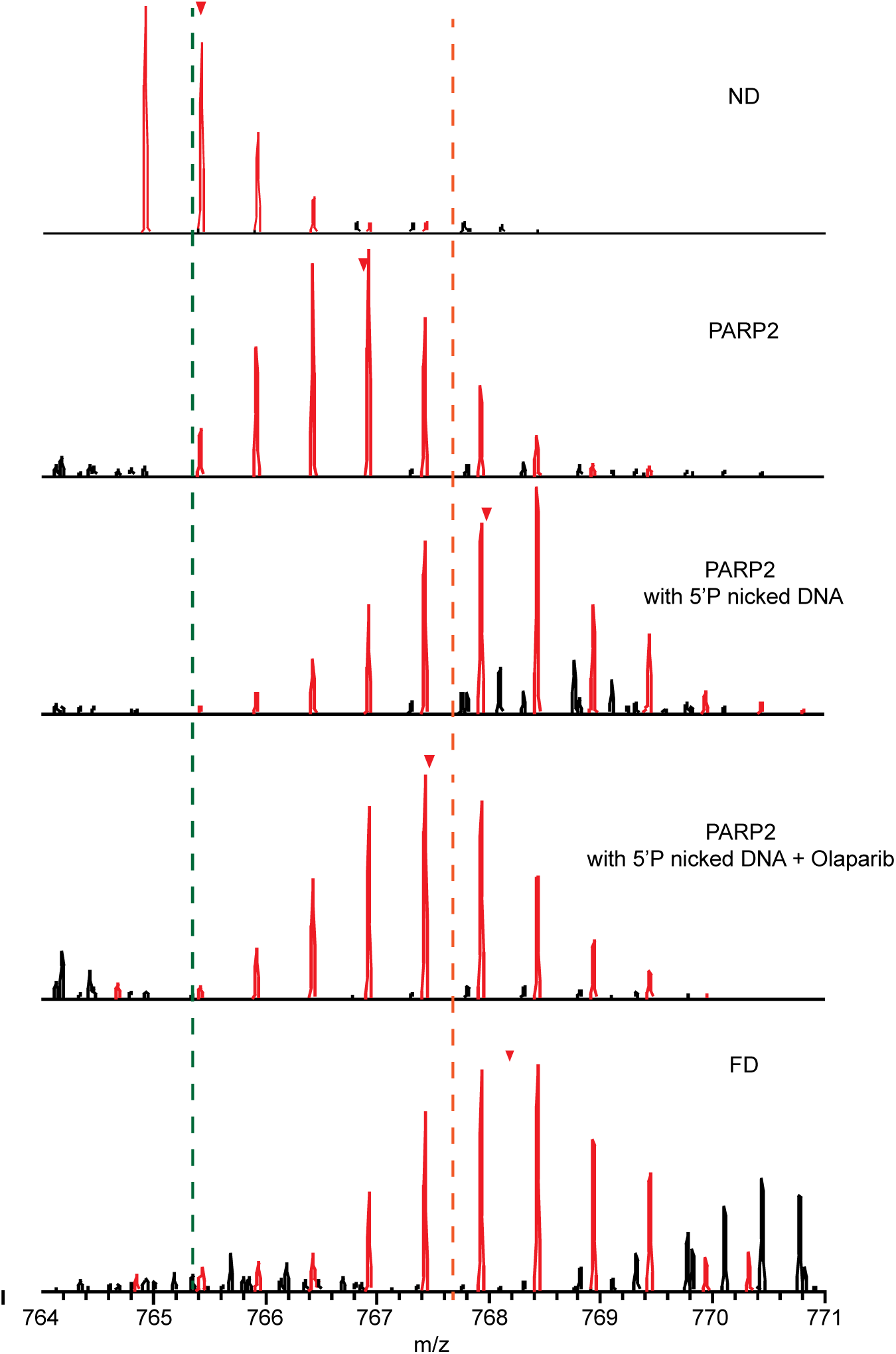
Raw MS data of representative peptide from ASL. Isotopic envelopes for the representative peptide of the ASL in an ND sample, PARP2, PARP2 with 5’P nicked DNA, PARP2 with 5’P nicked DNA and Olaparib, and FD sample at 100 s are shown. Red triangles indicate the centroid value. Green and orange dotted lines visualize the differences in m/z of the representative peptide of ASL.

## References

1. Pandey, N., and Black, B.E. (2021). Rapid detection and signaling of DNA damage by PARP-1. Trends Biochem Sci 46, 744–757. 10.1016/j.tibs.2021.01.014.

2. Ray Chaudhuri, A., and Nussenzweig, A. (2017). The multifaceted roles of PARP1 in DNA repair and chromatin remodelling. Nat Rev Mol Cell Biol 18, 610–621. 10.1038/nrm.2017.53.

3. Satoh, M.S., and Lindahl, T. (1992). Role of poly(ADP-ribose) formation in DNA repair. Nature 356, 356–358. 10.1038/356356a0.

4. Bonfiglio, J.J., Fontana, P., Zhang, Q., Colby, T., Gibbs-Seymour, I., Atanassov, I., Bartlett, E., Zaja, R., Ahel, I., and Matic, I. (2017). Serine ADP-ribosylation depends on HPF1. Mol Cell 65, 932–940.e6. 10.1016/j.molcel.2017.01.003.

5. Palazzo, L., Leidecker, O., Prokhorova, E., Dauben, H., Matic, I., and Ahel, I. (2018). Serine is the major residue for ADP-ribosylation upon DNA damage. eLife 7, e34334. 10.7554/eLife.34334.

6. Chen, Q., Kassab, M.A., Dantzer, F., and Yu, X. (2018). PARP2 mediates branched poly ADP-ribosylation in response to DNA damage. Nat Commun 9, 3233. 10.1038/s41467-018-05588-5.

7. Dawicki-McKenna, J.M., Langelier, M.-F., DeNizio, J.E., Riccio, A.A., Cao, C.D., Karch, K.R., McCauley, M., Steffen, J.D., Black, B.E., and Pascal, J.M. (2015). PARP-1 activation requires local unfolding of an autoinhibitory domain. Mol Cell 60, 755–768. 10.1016/j.molcel.2015.10.013.

8. Gibbs-Seymour, I., Fontana, P., Rack, J.G.M., and Ahel, I. (2016). HPF1/C4orf27 Is a PARP-1-Interacting Protein that Regulates PARP-1 ADP-ribosylation Activity. Mol Cell 62, 432–442. 10.1016/j.molcel.2016.03.008.

9. Langelier, M.-F., Billur, R., Sverzhinsky, A., Black, B.E., and Pascal, J.M. (2021). HPF1 dynamically controls the PARP1/2 balance between initiating and elongating ADP-ribose modifications. Nat Commun 12, 6675. 10.1038/s41467-021-27043-8.

10. Murai, J., Huang, S.N., Das, B.B., Renaud, A., Zhang, Y., Doroshow, J.H., Ji, J., Takeda, S., and Pommier, Y. (2012). Differential trapping of PARP1 and PARP2 by clinical PARP inhibitors. Cancer Res 72, 5588–5599. 10.1158/0008-5472.CAN-12-2753.

11. Ménissier de Murcia, J., Ricoul, M., Tartier, L., Niedergang, C., Huber, A., Dantzer, F., Schreiber, V., Amé, J.-C., Dierich, A., LeMeur, M., et al. (2003). Functional interaction between PARP-1 and PARP-2 in chromosome stability and embryonic development in mouse. EMBO J 22, 2255–2263. 10.1093/emboj/cdg206.

12. Langelier, M.-F., Riccio, A.A., and Pascal, J.M. (2014). PARP-2 and PARP-3 are selectively activated by 5′ phosphorylated DNA breaks through an allosteric regulatory mechanism shared with PARP-1. Nucleic Acids Res 42, 7762–7775. 10.1093/nar/gku474.

13. Riccio, A.A., Cingolani, G., and Pascal, J.M. (2016). PARP-2 domain requirements for DNA damage-dependent activation and localization to sites of DNA damage. Nucleic Acids Res 44, 1691–1702. 10.1093/nar/gkv1376.

14. Mortusewicz, O., Amé, J.-C., Schreiber, V., and Leonhardt, H. (2007). Feedback-regulated poly(ADP-ribosyl)ation by PARP-1 is required for rapid response to DNA damage in living cells. Nucleic Acids Res 35, 7665–7675. 10.1093/nar/gkm933.

15. Zandarashvili, L., Langelier, M.-F., Velagapudi, U.K., Hancock, M.A., Steffen, J.D., Billur, R., Hannan, Z.M., Wicks, A.J., Krastev, D.B., Pettitt, S.J., et al. (2020). Structural basis for allosteric PARP-1 retention on DNA breaks. Science 368, eaax6367. 10.1126/science.aax6367.

16. Langelier, M.-F., Lin, X., Zha, S., and Pascal, J.M. (2023). Clinical PARP inhibitors allosterically induce PARP2 retention on DNA. Sci Adv 9, eadf7175. 10.1126/sciadv.adf7175.

17. Lin, X., Jiang, W., Rudolph, J., Lee, B.J., Luger, K., and Zha, S. (2022). PARP inhibitors trap PARP2 and alter the mode of recruitment of PARP2 at DNA damage sites. Nucleic Acids Res 50, 3958–3973. 10.1093/nar/gkac188.

18. Englander, S.W. (2006). Hydrogen exchange and mass spectrometry: A historical perspective. J Am Soc Mass Spectrom 17, 1481–1489. 10.1016/j.jasms.2006.06.006.

19. Langelier, M.-F., Zandarashvili, L., Aguiar, P.M., Black, B.E., and Pascal, J.M. (2018). NAD^+^ analog reveals PARP-1 substrate-blocking mechanism and allosteric communication from catalytic center to DNA-binding domains. Nat Commun 9, 844. 10.1038/s41467-018-03234- 8.

20. Bilokapic, S., Suskiewicz, M.J., Ahel, I., and Halic, M. (2020). Bridging of DNA breaks activates PARP2–HPF1 to modify chromatin. Nature 585, 609–613. 10.1038/s41586-020-2725-7.

21. Obaji, E., Maksimainen, M.M., Galera-Prat, A., and Lehtiö, L. (2021). Activation of PARP2/ARTD2 by DNA damage induces conformational changes relieving enzyme autoinhibition. Nat Commun 12, 3479. 10.1038/s41467-021-23800-x.

22. Langelier, M.-F., Planck, J.L., Roy, S., and Pascal, J.M. (2012). Structural basis for DNA damage–dependent Poly(ADP-ribosyl)ation by human PARP-1. Science 336, 728–732. 10.1126/science.1216338.

23. Oliver, A.W., Amé, J., Roe, S.M., Good, V., de Murcia, G., and Pearl, L.H. (2004). Crystal structure of the catalytic fragment of murine poly(ADP-ribose) polymerase-2. Nucleic Acids Res 32, 456–464. 10.1093/nar/gkh215.

24. Karlberg, T., Hammarström, M., Schütz, P., Svensson, L., and Schüler, H. (2010). Crystal structure of the catalytic domain of human PARP2 in complex with PARP inhibitor ABT-888. Biochemistry 49, 1056–1058. 10.1021/bi902079y.

25. Thorsell, A.-G., Ekblad, T., Karlberg, T., Löw, M., Pinto, A.F., Trésaugues, L., Moche, M., Cohen, M.S., and Schüler, H. (2017). Structural basis for potency and promiscuity in Poly(ADP-ribose) Polymerase (PARP) and Tankyrase inhibitors. J Med Chem 60, 1262–1271. 10.1021/acs.jmedchem.6b00990.

26. Papeo, G., Posteri, H., Borghi, D., Busel, A.A., Caprera, F., Casale, E., Ciomei, M., Cirla, A., Corti, E., D’Anello, M., et al. (2015). Discovery of 2-[1-(4,4-Difluorocyclohexyl)piperidin-4-yl]- 6-fluoro-3-oxo-2,3-dihydro-1H-isoindole-4-carboxamide (NMS-P118): a potent, orally available, and highly selective PARP-1 inhibitor for cancer therapy. J Med Chem 58, 6875–6898. 10.1021/acs.jmedchem.5b00680.

27. Suskiewicz, M.J., Zobel, F., Ogden, T.E.H., Fontana, P., Ariza, A., Yang, J.-C., Zhu, K., Bracken, L., Hawthorne, W.J., Ahel, D., et al. (2020). HPF1 completes the PARP active site for DNA damage-induced ADP-ribosylation. Nature 579, 598–602. 10.1038/s41586-020-2013-6.

28. Yu, H., Ma, H., Yin, M., and Wei, Q. (2012). Association between PARP-1 V762A polymorphism and cancer susceptibility: a meta-analysis. Genet Epidemiol 36, 56–65. 10.1002/gepi.20663.

29. Wang, X.-G., Wang, Z.-Q., Tong, W.-M., and Shen, Y. (2007). PARP1 Val762Ala polymorphism reduces enzymatic activity. Biochem Biophys Res Commun 354, 122–126. 10.1016/j.bbrc.2006.12.162.

30. Gui, B., Gui, F., Takai, T., Feng, C., Bai, X., Fazli, L., Dong, X., Liu, S., Zhang, X., Zhang, W., et al. (2019). Selective targeting of PARP-2 inhibits androgen receptor signaling and prostate cancer growth through disruption of FOXA1 function. PNAS 116, 14573–14582. 10.1073/pnas.1908547116.

31. Galindo-Campos, M.A., Lutfi, N., Bonnin, S., Martínez, C., Velasco-Hernandez, T., García-Hernández, V., Martín-Caballero, J., Ampurdanés, C., Gimeno, R., Colomo, L., et al. (2022). Distinct roles for PARP-1 and PARP-2 in c-Myc–driven B-cell lymphoma in mice. Blood 139, 228–239. 10.1182/blood.2021012805.

32. Steffen, J.D., Brody, J.R., Armen, R.S., and Pascal, J.M. (2013). Structural implications for selective targeting of PARPs. Front Oncol 3, 301. 10.3389/fonc.2013.00301.

33. Rolli, V., O’Farrell, M., Ménissier-de Murcia, J., and de Murcia, G. (1997). Random mutagenesis of the Poly(ADP-ribose) Polymerase catalytic domain reveals amino acids involved in polymer branching. Biochemistry 36, 12147–12154. 10.1021/bi971055p.

34. Rouleau-Turcotte, É., Krastev, D.B., Pettitt, S.J., Lord, C.J., and Pascal, J.M. (2022). Captured snapshots of PARP1 in the active state reveal the mechanics of PARP1 allostery. Mol Cell 82, 2939–2951.e5. 10.1016/j.molcel.2022.06.011.

35. Shao, Z., Lee, B.J., Zhang, H., Lin, X., Li, C., Jiang, W., Chirathivat, N., Gershik, S., Shen, M.M., Baer, R., et al. (2023). Inactive PARP1 causes embryonic lethality and genome instability in a dominant-negative manner. PNAS 120, e2301972120. 10.1073/pnas.2301972120.

36. Dréan, A., Lord, C.J., and Ashworth, A. (2016). PARP inhibitor combination therapy. Crit Rev Oncol Hematol 108, 73–85. 10.1016/j.critrevonc.2016.10.010.

37. Lord, C.J., and Ashworth, A. (2017). PARP inhibitors: Synthetic lethality in the clinic. Science 355, 1152–1158. 10.1126/science.aam7344.

38. Rose, M., Burgess, J.T., O’Byrne, K., Richard, D.J., and Bolderson, E. (2020). PARP inhibitors: clinical relevance, mechanisms of action and tumor resistance. Front Cell Dev Biol 8. 10.3389/fcell.2020.564601.

39. Illuzzi, G., Staniszewska, A.D., Gill, S.J., Pike, A., McWilliams, L., Critchlow, S.E., Cronin, A., Fawell, S., Hawthorne, G., Jamal, K., et al. (2022). Preclinical characterization of AZD5305, a next-generation, highly selective PARP1 inhibitor and trapper. Clin Cancer Res 28, 4724–4736. 10.1158/1078-0432.CCR-22-0301.

40. Papeo, G., Orsini, P., Avanzi, N.R., Borghi, D., Casale, E., Ciomei, M., Cirla, A., Desperati, V., Donati, D., Felder, E.R., et al. (2019). Discovery of stereospecific PARP-1 inhibitor isoindolinone NMS-P515. ACS Med Chem Lett 10, 534–538. 10.1021/acsmedchemlett.8b00569.

41. Velagapudi, U.K., Rouleau-Turcotte, É., Billur, R., Shao, X., Patil, M., Black, B.E., Pascal, J.M., and Talele, T.T. (2024). Novel modifications of PARP inhibitor veliparib increase PARP1 binding to DNA breaks. Biochem J 481, 437–460. 10.1042/BCJ20230406.

42. Langelier, M.-F., Steffen, J.D., Riccio, A.A., McCauley, M., and Pascal, J.M. (2017). Purification of DNA damage-dependent PARPs from E. coli for structural and biochemical analysis. Methods Mol Biol 1608, 431–444. 10.1007/978-1-4939-6993-7_27.

